# Scaly-Tail Organ Enhances Static Stability during Pel’s Scaly-tailed Flying Squirrels’ Arboreal Locomotion

**DOI:** 10.1101/2024.12.31.630952

**Authors:** Andrew K. Schulz, Mrudul Chellapurath, Pranav C. Khandelwal, SeyedReza Rezaei, Stefan Merker, Ardian Jusufi

## Abstract

Scaly-tailed squirrels (*Anomaluridae*) are one of the least studied mammalian families. Their namesake is due to a peculiar and unique scaly-tail organ extruding from the caudal vertebrate that has been predicted to help reduce skidding. This study investigates the function of the scaly-tail organ found in *Anomalurus pelii*, investigating its potential role in enhancing arboreal locomotion. As these animals glide from tree to tree in a habitat abundant with smooth-bark trees, we hypothesize that the scaly-tail organ assists with friction enhancement in their native smooth-bark habitat. Through a combination of analyses using mathematical and physical models for experimental validation, we explore whether the scaly-tail organ could improve the sliding and overturning stability during perching. Our experimental results showed that the scaly-tail organ can act as a skid-reduction mechanism by enhancing substrate engagement on intermediate roughness substrates by 58%. Mathematical models showed the scaly-tail organ enhances overturning stability by acting as an additional support point. Our model showed that the scaly-tailed squirrel can reach up to 82.5^◦^ inclination without claw force; however, without scales, it reduces to 79.6^◦^. Overall, this research highlights the functional significance of scaly-tail organs in adaptations in scaly-tailed flying squirrels and contributes to our understanding of their locomotion strategies and environmental stresses. Our study also provides insights into innovative perching mechanisms for robots operating in arboreal environments.

## 1. Introduction

Arboreal environments are highly three-dimensional discontinuous structures in which animals move horizontally and vertically. They do so by performing behaviors including climbing, jumping, perching, hanging, descending, and even gliding [1] while interacting with various substrates, from rough tree barks or ranches to relatively smooth, slippery surfaces. The variability of surface texture and composition necessitates different behavioral and morphological adaptations to maintain traction and stability and not fall while performing these behaviors [2]. Large body-sized animals like primates use long appendages to grasp and hold tree branches and bridge gaps [3]. Snakes use their elongated bodies to generate sufficient muscular force to wrap and grip the arboreal surface [4]. As body size decreases, morphological adaptations such as prehensile tails, claws, spines, or adhesive pads become more prevalent [5]. These adaptations are seen in animals spanning approximately seven orders of magnitude in body mass, from mites to lizards [6], [7], [8].

Morphological adaptations such as claws and adhesive pads have been extensively studied in lizards and have shown to be an effective way to grip the arboreal substrate and generate substrate reaction forces during climbing [9]. The insights from these studies have also been applied to bio-inspired robots, enabling them to scale inclines and vertical substrates [10]. Additionally, claws and adhesive pads [11], can be complemented by the use of the tail, which can act as a fifth point of contact between the substrate and the body. For example, chameleons exhibit a prehensile tail, which can be coiled around a perch during slow climbing to act as an anchor point while crossing gaps [12]. Also the distal tail tips of birds can be used to support them during climbing, which is achieved through geometry and not material shifts [13]. Treecreepers can use the tail tip anchorage with specialized feathers for station holding during large impulse dissipation as the beak causes perforation in the tree trunk [14].

When it comes to dynamic vertical locomotion, geckos were found to use a tail reflex to reject perturbations, with reduced climbing performance documented in tailless animals [15]. Further substantiating the hypothesis of the function of contact tails in disturbance rejection during rapid vertical running, the tails of highly arboreal lizards were seen to be in contact during climbing of rough substrates [16]. It was hypothesized to be aided by morphological features on the ventral side of the tail, keeled subcaudal scales with spines pointing distally, the forces individual scales could sustain were experimentally found to support body mass and beyond in three species [16].

Gliding geckos can press their tail against the substrate to prevent pitch back and falling during landing on trees [16], [17]. Active tail movement can also facilitate mid-air torso reorientation by appendage inertia, as has been seen in lizards and squirrels [15], [18], [19], [20]. The role of the active tail robot’s [17] back and tail stiffness was evaluated with at-scale physical models of perching geckos [21]. All of these studies highlight the role of the tail as an appendage with functions critical to locomotion performance. Interestingly, as seen in highly arboreal lizards [16], the tails of some arboreal animals also possess specialized morphological features such as spines and scales, which can potentially provide additional mechanical support during climbing or perching [22]. However, their role remains under-explored. Robot locomotion experiments demonstrate that by combining these biological systems with engineering principles, such as shape-morphing [23] or variable stiffness [24], additional benefits can be achieved, including generalist locomotion or reduction of energy cost.

In this study, we investigate the role of one such tail adaptation found in the elusive scaly-tailed squirrels, a family of rodents native to Africa. These squirrels have a unique tail adaptation called the ‘scaly-tail organ’ with scales that protrude from the caudal spine bones out of the skin (**Figure 1**, **Figure S1**) [22]. This adaptation is shared across all species in the family *Anomaluridae*, which include *Anomalurus pusillus*, *A. beecrofti*, *A. derbianus*, and *A. pelii* [22]. Moreover, the *Anomaluridae* species use gliding to traverse their habitat (but see *Zenkerella insignis* [23]). Still, they are more closely related to non-gliding rodents than they are to flying squirrels, which belong to the family *Sciuridae* (**Figure 1, Table S1**). A distinctive feature of the Scaly tailed squirrels’ habitat, specifically that of *Anomalurus pelii* or Pel’s scaly-tailed squirrel, is a high abundance of trees that are often referred to as ‘smooth-barked’ trees [24] shown in **Figure 1**D. These trees have barks that are much smoother than tree barks found in habitats of flying squirrels that lack the scaly tail adaptation, such as the Mentawi flying squirrel (*Iomys sipora*), Woolly Flying Squirrel (*Eupetaurus cinereus*), and Sipora flying squirrel (*Hylopetes sipora*)[25]. The unique tail adaptation in the Pel’s scaly-tailed squirrel habitat has led to the hypothesis that the squirrel’s scaly-tail organ has evolved as a skid reduction mechanism and to support body weight while perching on the tree trunk [26], [27]. However, these hypotheses are derived from field observations and have not been supported by experiments [28], [29], [30], [31].

**Figure 1:**
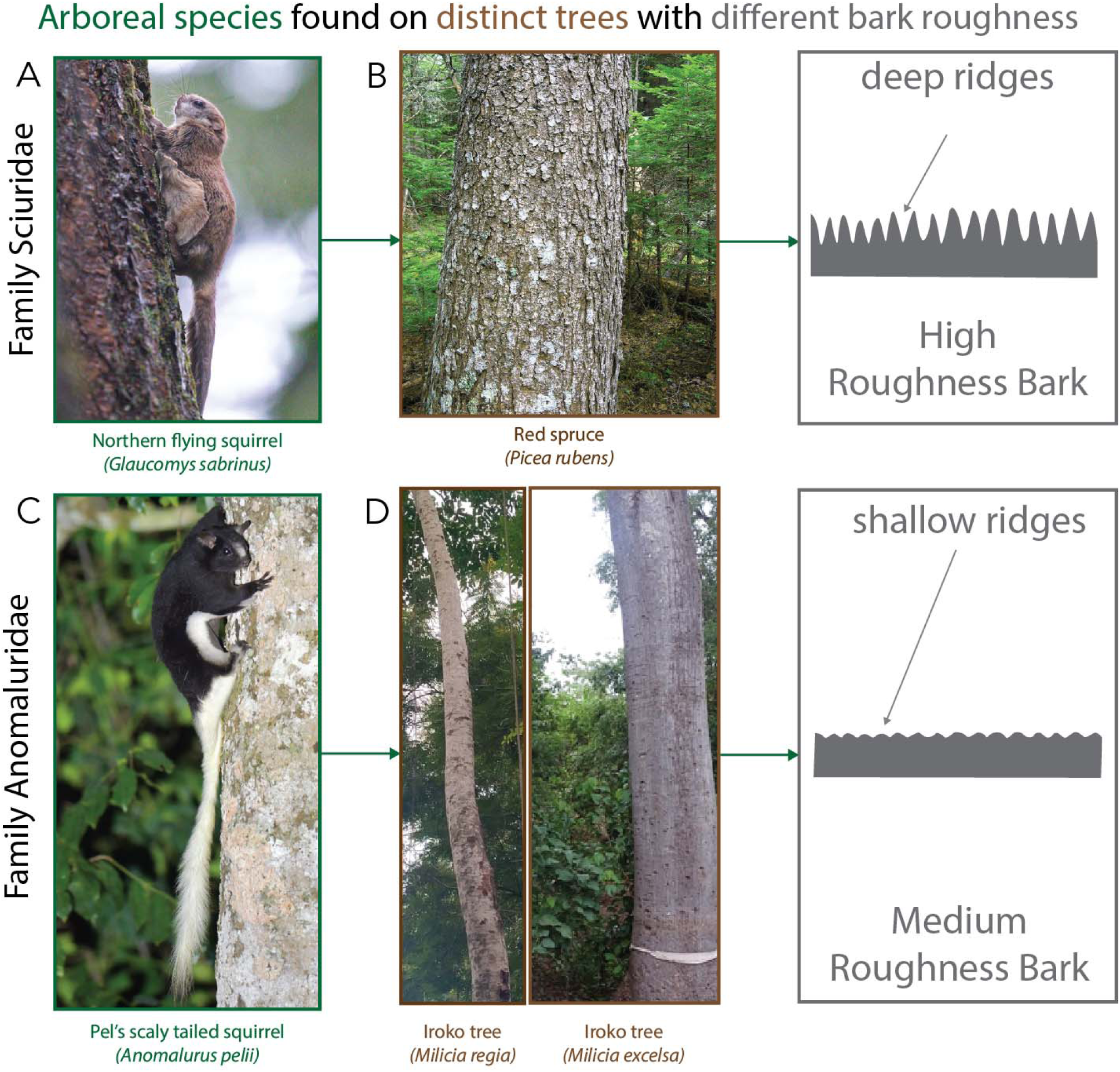
Arboreal mammalian species with native trees used for climbing. A) Image of Northern Flying Squirrel (*Glaucomys sabrinus*) climbing a tree photo taken by photospaul. B) Image of red spruce, commonly climbed by the northern flying squirrel, with a schematic showing the deep ridges from the scaled surface of the spruce tree. C) Image of the Pel’s scaly tailed squirrel (*Anomalurus pelii*) we examine in this study taken by pfaucher D) two common trees found in their native habitat of West Africa, the *Melicia regia* and *Milicia excelsa*. As the schematic shows, these are described as smooth bark trees with smooth surfaces and small ridges caused by creases along the tree. Photos taken by agboola and Sadamtoro. Photos are taken from iNaturalist, all under creative license CC BY-NC.

**Figure 2:**
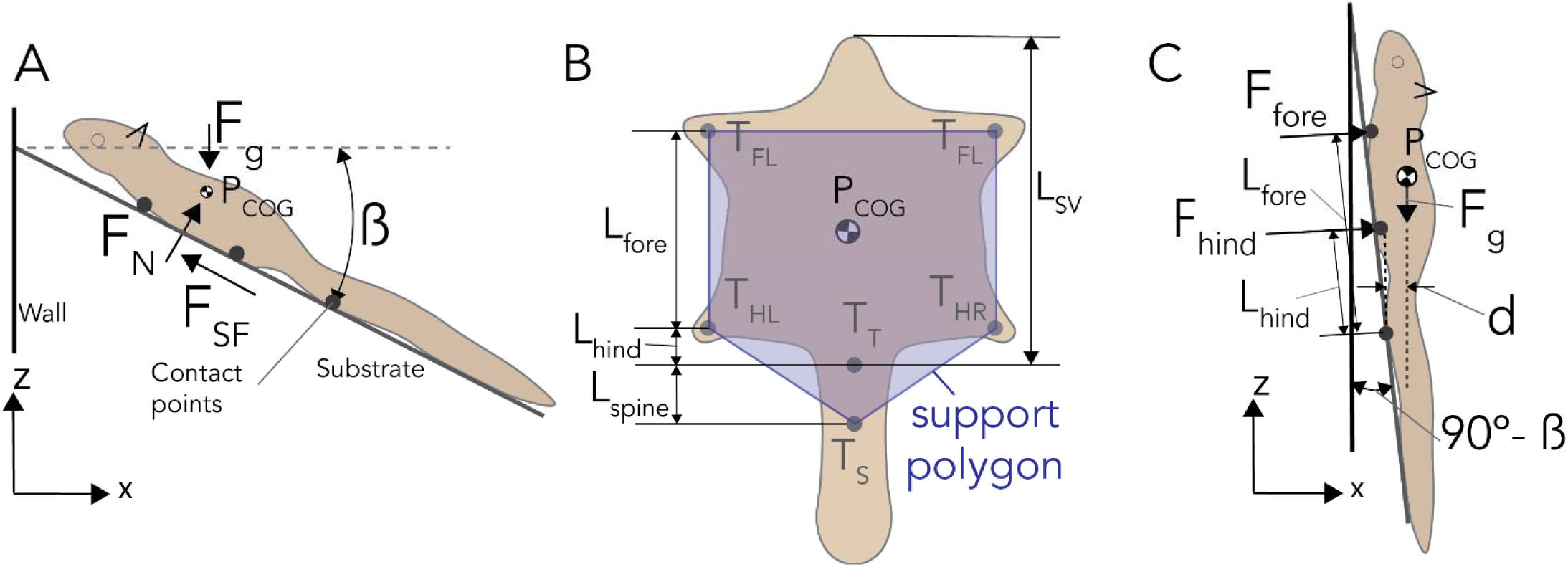
Stability of squirrel on an inclined surface. A) Forces acting on the squirrel resting on an inclined surface, B) Support polygon formed by the four claws and the scales of the tail, and the C) Force applied by the claws to resist the overturning due to the moment by the gravitational force.

Our study addresses these hypotheses and further investigates the scaly-tail organ’s role in enhancing the squirrel’s perching capabilities. We do this by characterizing the three-dimensional (3D) shape of the Pel’s scaly-tailed squirrel claw and scaly-tail organ, and using that to perform a static stability analysis of the squirrel’s perching behavior. We analyze two types of static instability that any animal/object can encounter on an inclined surface: sliding and overturning. Sliding happens when the component of gravitational force is greater than the frictional resistance provided by the points of contact with the substrate. Overturning can take place when the horizontal projection of the squirrel’s Centre of Gravity (COG) moves out of the support polygon (defined as the convex polygon formed by connecting the supports on the substrate) [32].

Using a 3D-printed physical model of the scaly-tail organ and claws, we investigated the sliding stability with experiments on an angular varying sandpaper ramp. We proceeded with mathematical simulations to test the impact of the scaly-tail organ on overturning stability during perching. Overall, the sliding stability analysis tested the hypothesis that the scaly-tail organ acts as a skid reduction mechanism for the squirrel and can support body weight on ‘smooth’ bark-like surfaces compared to the squirrel model without the scaly-tail organ. Specifically, we predicted that the scaly-tail organ would show similar frictional characteristics as the claws on extremely smooth substrates but significantly improve substrate engagement on intermediate roughness substrates. Furthermore, we hypothesized that the scaly-tail organ size is adapted to maximize the perch angle, and a smaller or larger scaly-tail organ size would reduce the perch angle, likely due to poorer engagement of the scales with the substrate.

The overturning instability was investigated using a mathematical model that simulated different scaly-tail organ sizes and the squirrel’s COG position to understand how the support polygon formed by the fore-hindlimb claws and the scaly-tail organ influenced overturning during perching. Based on this analysis approach, we hypothesized that the scaly-tailed organ enhances the over-turning stability of the squirrel while perching by increasing the area of the support polygon within which the projection of COG lies compared to a support polygon formed by the fore and hindlimb claws only. The presence of spines are critical in this case because at steeper angles of perching, the tail without the spines would slip and will not be able to act as a support point. Specifically, we would observe a steeper incline threshold for overturning for support polygons formed with the scaly-tailed organ compared to the polygon with only four claws.

## 2. Materials and Methods

### 2.1 Squirrel morphological data

Morphological data were taken from a single specimen of Pel’s scaly-tailed squirrel, also known as Pel’s scaly squirrel (*Anomalurus pelii*), at the State Museum of Natural History Stuttgart (SMNS) in November 2022. The specimen skin (SMNS-Z-MAM-001377) is shown in **Figure 3**A, which was collected in Ghana in 1881.

**Figure 3:**
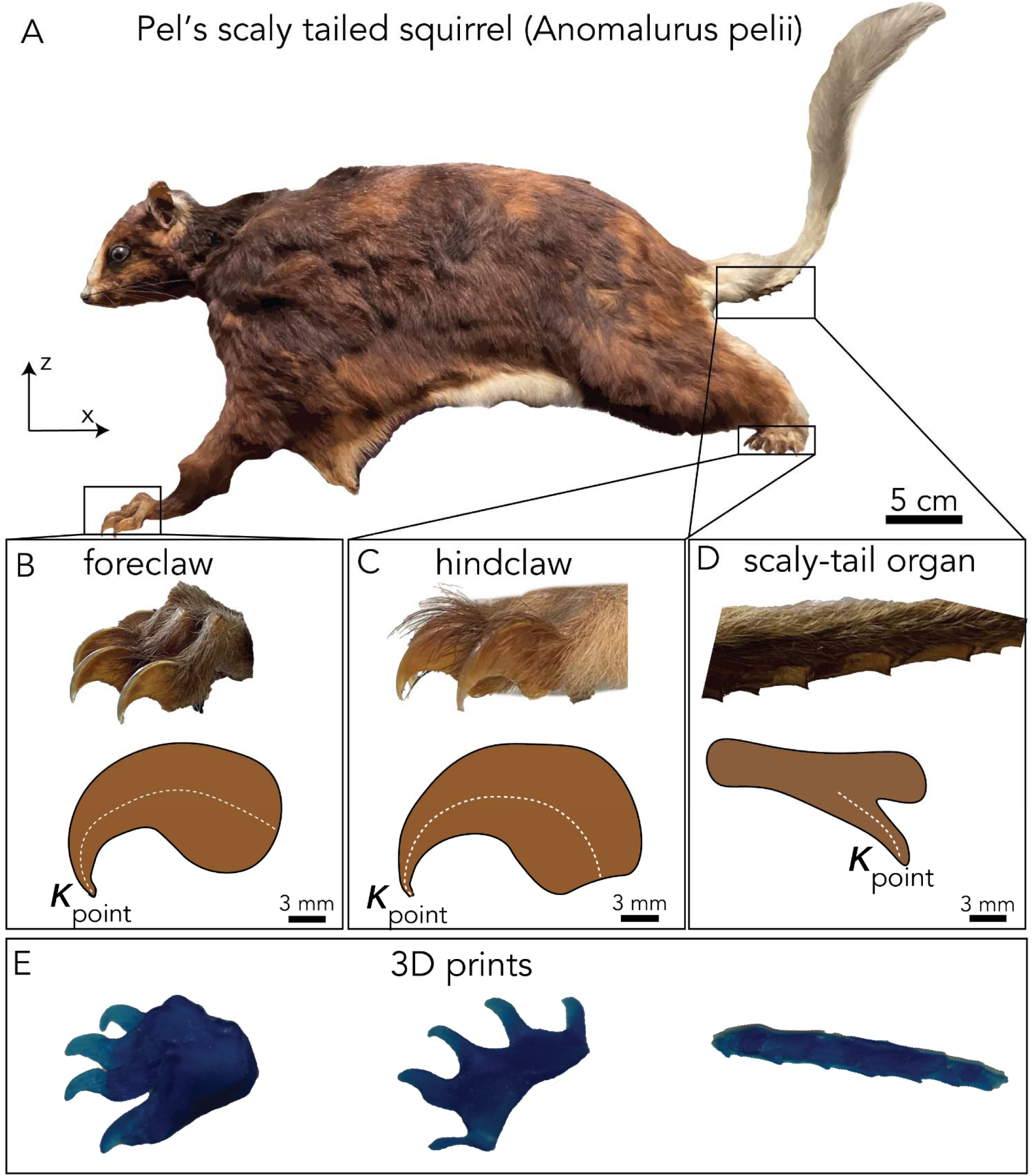
3D scanned portions of Pel’s scaly-tailed. squirrel A) Mounted museum specimen (SMNS-Z-MAM-001377) of *Anomalurus pelii* used for 3D scanning, B-D) foreclaw, hind claw, and scaly organ of the squirrel and a side view of the curvature and point curvature of each keratinized section of the squirrel. E) 3D printed foreclaw, hindclaw, and scaly organ from scans of squirrel displayed.

Photographs of the entire squirrel and close-ups of the claws and spiny tail were taken against a 1x1 cm graph sheet as shown in **Figure 3**A-D and **Figure S2**. Post photographs, the mounted Pel’s scaly-tailed squirrel was 3D scanned using a handheld scanner (Artec Space Spyder 3D Scanner) by placing it on a plastic turntable. Scanning was performed at eight frames per second (fps) at a distance ranging from 18 to 30 cm away from the specimen. Six scans were performed, including two scans of each site: the scaly-tail organ, foreclaw, and the hindclaw. Due to the staging of the mounted specimen in the museum collection, the right forepaw and left hindpaw were partially obstructed and could not be scanned. Therefore, the other hindclaw and foreclaw were mirrored to complete the scanned results.

The Artec 3D scans traced both the geometry and texture. The total scanned surfaces differed between the foreclaw, hindclaw, and the tail organ, as the surface size varies based on the sample volume scanned. Scanned samples averaged 800 surfaces for the claws and 1200 for the caudal scaly-tail organ.

The scans taken of each structure were imported into Artec 3D Professional from Artec Studio 18, and the raw scans were cropped only to include the structures of interest (i.e., claws and scaly-tail organ). The scans were converted into 3D mesh structures using the Artec software’s automated tool. The minor irregularities in the 3D model, like small holes, were manually filled and fitted with the default spline interpolation provided in the software. Moreover, smoothing allowed removing any spikes in the scan caused by the hair on the sample, leading to a solid structure for 3D printing (**Figure 3**E).

Finally, the 3D scans were used to take measurements of the foreclaw, hindclaw, and scaly-tailed organ (**Figure 3**B-D). The photographs were used to measure the body and appendage length reported in the supplement (**Figure S3, Table S3**).

The curvature of the claws/scale tips was taken as the two-dimensional curvature based on an arclength of the line surrounding the structure. We denote this arclength function as *y*(*x*), which is the function of a position, *x*. We can find the curvature of this arclength by using the function in the curvature equation given by:

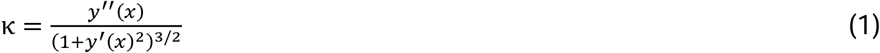

where *y*(*x*) is a function of the center-line of curvature, we take the curvature along the tips of each of the claws/spines to calculate the ball tip curvature as seen in **Figure 3**B-D.

### 2.2 Sliding stability analysis

#### 2.2.1 Squirrel physical model creation

The solid 3D models generated using the Artec software were printed (**Figure 3**E) on a Stratasys J835 PolyJet 3D printer using Vero transparent blue printing material. This material has an approximate tensile strength of 50 to 60 MPa and a modulus of elasticity of 2000 to 3000 MPa [33].

To create a physical model for frictional tests, we geometrically scaled down all morphological data of the Pel’s scaly tailed squirrel to half the size of the scanned squirrel (**Table S1, Figure S2**). The model’s mass was proportionately scaled down using the scaling law (*m* ∝ *l*^3^), which made the entirety of the physical model approximately 0.5 kg.

The scaled down appendage and body lengths of the museum specimen were used to create an approximate body structure. A simple skeleton was designed in SOLIDWORKS 2022 and fabricated via a laser cutting machine (Universal Laser System PLS6.150D). The foreclaws, hindclaws, and the scaly-tail organ were attached to the body structure as shown in **Figure 4**A-B. Finally, four black color ball pins were placed on the dorsal side of the model to enable image tracking.

**Figure 4:**
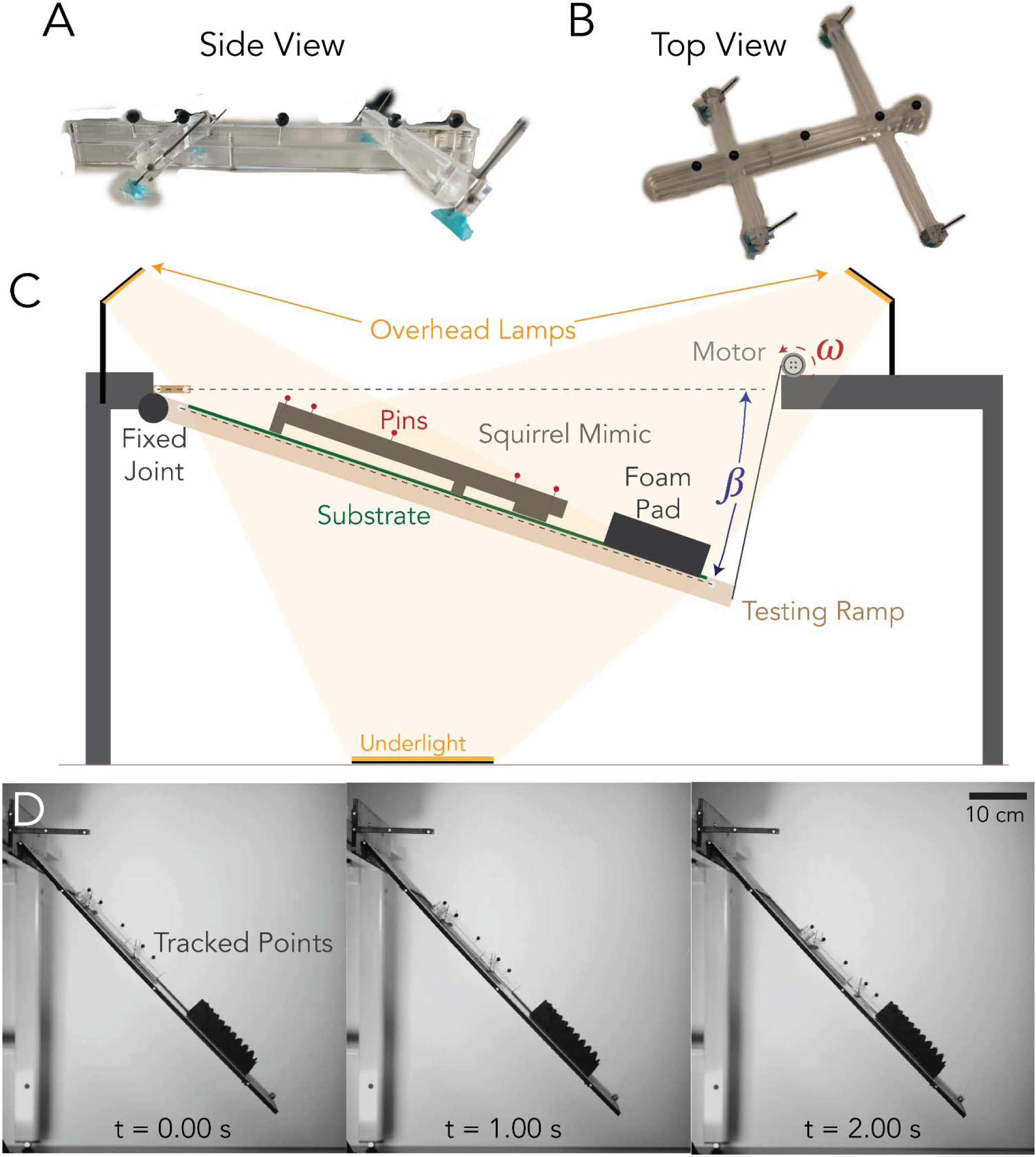
A) Top view of scaly squirrel mimic displaying forelimb and hindlimb of the Pel’s scaly tailed squirrel morphology, B) side view displaying the foreclaw, hindclaw, and scaly tail organ interacting with the surface. Overall frictional setup displaying the mimic descending with motor spinning at speed ω to give angle β from the horizontal axis. D) Labeled data of squirrel mimic on the setup showing time series of the experimental setup of the squirrel mimic with scales attached on P1500 sandpaper (12.6 ± 1 *µ*m average surface roughness) showing the movement and contact with the foam pad.

#### 2.2.2 Experimental setup

The role of the scaly-tail organ in the squirrel’s sliding stability was characterized through two experiments. The first experiment tested the frictional contribution of the scaly-tail organ compared to the claws using two versions of the squirrel physical model. Version one consisted of the model with all four claws and the scaly-tail organ; the second version removed the scaly-tail organ. Each version was tested on four different substrates represented by sandpapers of grit size P1500 (12.6±1 *µ*m), P600 (25.8±1 *µ*m), P150 (90±15 *µ*m), and P60 (270±10 *µ*m) [34]. A higher number following ‘P’ represents a ‘smoother’ grit paper and the value reported in the bracket is the root-mean-square average roughness (*Rq*) in *µm*. The *Rq* value allowed the sandpapers to be compared to tree barks of varying roughness. Previously published literature indicates that bamboo is similar to P400 sandpaper and P40 sandpaper is similar to high-roughness tree bark [35]. Therefore, we chose sandpapers that represent barks smoother than bamboo and less rough (intermediate) than high roughness bark trees like the red spruce which is inhabited by the northern flying squirrel [36]. Such a gradation allowed us to investigate the roughness at which the scaly-tail organ engages and potentially fails.

The second experiment tested the effect of the scaly-tail organ size on substrate engagement. Instead of the squirrel model, we used a rectangular plate mounted with the scaly-tailed organ. We tested five sizes of the scaly-tail organ by scaling it by the following ratios: 0.5x, 0.75x, x, 1.25x, and 1.5x. These ratios resulted in scale sizes of ball tip radii of 1.28, 1.92, 2.56, 3.20, and 3.84 mm, respectively. These sizes of the scaly-tail organ were tested on a single substrate (P60), which corresponded to the substrate that resulted in the steepest perch angle that could be sustained by the squirrel physical model during the sliding experiment.

Both experiments used the same setup consisting of a ramp with changing inclination, a sub-strate material, and an orthogonally placed monochrome high-speed camera (AOS-Smotion-104-M with AOS 50 mm lens) (**Figure 4**C-E). At least ten trials for each iteration of the experiment were performed, out of which some were discarded due to issues with the camera trigger and tracking (**Table S3**).

##### Ramp setup

The ramp was constructed using plywood (40 cm x 70 cm x 1 cm) with a placeholder (30 cm x 50 cm) in the middle for substrate placement (**Figure 4**C). One end of the ramp was connected to a hinge on an elevated platform, and the other end was connected to a pulley through a cable. The inclination of the ramp was controlled using a servomotor (Dynamixel WC430-W150-T) connected to the pulley mechanism **Figure 4**C. The inclination was varied from a horizontal position down to a maximum of -90 ^◦^ at a speed of 0.229 rpm. Four white color ball pins were placed along the side (thick edge) of the ramp to provide high-contrast points for motion tracking. An L-shaped plywood piece with four white color ball pins was placed on the top left of the ramp such that the longer side of the L-shaped piece was horizontal with the ground.

##### Experimental protocol

For each experiment, the substrate was mounted on the ramp, followed by placing the model on the substrate. The topmost point of contact of the model was near the top edge of the substrate, providing sufficient space on the ramp for it to slide freely. The experiment started from a ramp inclination of 0^◦^ and increased at a rate of -0.94 ± 0.42 degrees/s when averaged over all trials used for analysis (n = 123). Simultaneously, the high-speed camera recorded the trial at 200 fps with a 2 s before and after trigger buffer. As soon as the model slipped, the camera was triggered, allowing for the precise measurement of the ramp angle at which sliding occurred along with the velocity and acceleration of the model during each slide.

##### Data processing

The slip angle of the model was calculated by tracking the ball pins on the physical model/rectangular plate, ramp, and the L-shaped plywood piece using the deep learning pose estimation package DeepLabCut (DLC) [37]. A Resnet50 network was trained on 40 images of the scaly squirrel model, and a second Resnet50 network was trained on 201 images of the rectangular plate with the scaly-tail organ (**Figure 4**D). The tracking output from the DLC package consisted of the pixel coordinates of all the ball pins for the entire video duration. The pixel coordinates were calibrated using the actual measurement between two ball pins on the horizontal leg of the L-shaped piece. After calibration, each ball pin track on the squirrel model, rectangular plate, and ramp was smoothed using a smoothing quintic spline, followed by taking the smoothed tracks’ first and second derivatives to calculate the 2D velocity and acceleration. Each track was translated and rotated such that the origin (X = 0, Y = 0) corresponded to the pin on the ramp closest to the hinge and the positive X axis. The instance when the model slipped was identified by the sudden change in the distance of the ball pin on the model with respect to the origin. The sudden change corresponded to a change in the slope of the distance curve plotted with time. The frame at which slip was detected was used to calculate the ramp angle and corresponded to the model’s slip angle for the static friction calculation (*µ_s_*).

### 2.3 Coefficient of static friction calculation

Here, we derive the coefficient of static friction equation for the squirrel model, and the same equation applies to the rectangular plate with the scaly-tail organ. The forces acting on the squirrel when it is statically stable are the gravitational force *F_G_*, Normal force *F_N_*, and the Static frictional force *F_SF_* (**Figure 2A**). We assume the force of static friction increases linearly with the applied force until the maximum value is reached. Therefore, we know that the static friction force is

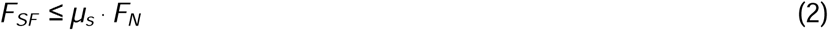

where *µ_s_* is the static friction coefficient between the squirrel and the tree. This value is specific to the material differences between the squirrel and the tree. Applying the first condition for equilibrium,

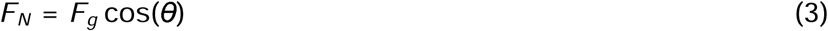

and

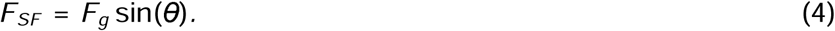

Substituting the value of *F_N_* and *F_SF_* in Equation (2), we simplify to:

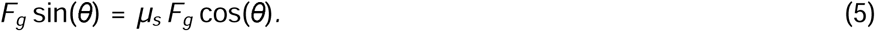

Finally, we can solve this equation for the static friction coefficient *µ_s_*:

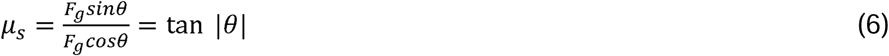

Where *θ*, in this case, is the inclination of the substrate. The maximum value of *µ_s_* is tan *θ_max_*, where *θ_max_* is the angle at which the squirrel model starts sliding.

### 2.4 Overturning stability analysis

The utility of the tail during landing has been observed in gliding vertebrates landing on vertical tree trunks with dynamic model showing that forces required at the foot would be inversely proportional to length of tail [16]. Our study focused on the static stability and hence based our model on the concept of support polygon [38]. The squirrel gets overturned when the Centre of gravity (COG) projection falls outside the support polygon. In this case, the support polygon is the convex polygon formed by connecting the attachment formed by the claws (*T_FL_*,*T_FR_*, *T_HL_*, and *T_HR_*) and the spiny tail (*T_S_*), as shown in **Figure 2**B. It is assumed that the claws and spiney tail are single support points and these points does not slide (**Figure 2**C).

When the squirrel is perching on an inclined surface, and the projection of COG (*P_COG_*) is inside the support polygon, the squirrel is statically stable. If the inclination is higher and the projection falls outside the support polygon (**Figure 2**C), the squirrel becomes statically unstable due to the moment generated by *F_COG_*. To avoid the pitch-back of the body about the tail base (*T_T_*), the squirrel has to actively apply force by using its claws to counter the moment generated by *F_COG_*. The position of *T_S_* (i.e., *L_spine_*) influences the maximum inclination angle of the substrate that the squirrel can stand stably without actively attaching to the substrate (i.e., the inclination angle at which the projection of COG falls on *T_S_*).

The maximum inclination till the squirrel is statically stable is given by:

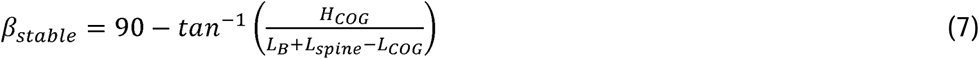

where, *H_COG_* is the height of COG, *L_COG_* is the distance of the COG from the snout, *L_B_* is the body length. We computed the value of β*_stable_* for a range of biologically possible values of *L_spine_*. The sum of moment force required to counter the moment by *F_COG_* is given by:

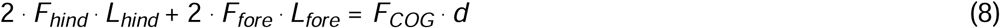

where *F_hind_* and *F_fore_* are the force produced by the hind and fore claws, respectively. The lengths, *L_hind_* and *L_fore_*, are the distance of hind and fore claws from the tail base, respectively, and *d* is the moment arm of *F_COG_*. We assume the forces produced by hind claws are 0.8 times the fore claw; (*F_fore_* = 0.8 *F_hind_*)[39], therefore ’Equation (8) can be simplified as:

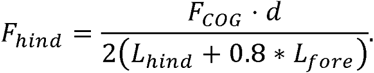

The moment arm, *d* can be calculated from the inclination angle, β, and morphological data of the squirrel:

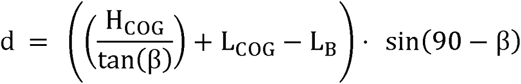

It is clear from this equation that the position of COG can influence the force the claws need to generate. To study the influence of the position of COG on the force required by the claws to prevent overturning, we measured the *F_claw_* for a range of *H_COG_* (*H_COG_*-2α, *H_COG_*-α, *H_COG_*, *H_COG_*+α, and *H_COG_*+2α) and *L_COG_* (*L_COG_*-2α, *L_COG_*-α, *L_COG_*, *L_COG_*+α, and *L_COG_*+2α), where α is 1 cm. For our analysis, we assume *P_COG_* is 4 cm away from the chest (*H_COG_*) and 17 cm from the head (*L_COG_*). We assume this because the total body diameter of the squirrel is 10 cm, and most mammals have a higher mass closer to their chest [40]. We assume the distance of the COG from the head is near the midway of the chest cavity but is slightly skewed towards the head as the head is more massive than the tail, which is primarily hair and fur. All reported numbers from the stability analysis are rounded to three significant digits to be consistent with the rest of the manuscript.

### 2.5 Statistical and data analysis

All calculations, including statistical analysis, were performed in MATLAB 2023a (The MathWorks, Natick, MA, USA) using custom scripts (included as supplementary material). The average metrics reported throughout this manuscript are in the form of mean ± std, and are averaged across all trials for each substrate (**Figure 5A-B**) or per spine size (**Figure 5**E). The test for significance between different groups was performed using the Wilcoxon rank sum test for unequal sample size. The morphological data used are taken from the measurements presented in Table S3 in Supplementary Material.

**Figure 5:**
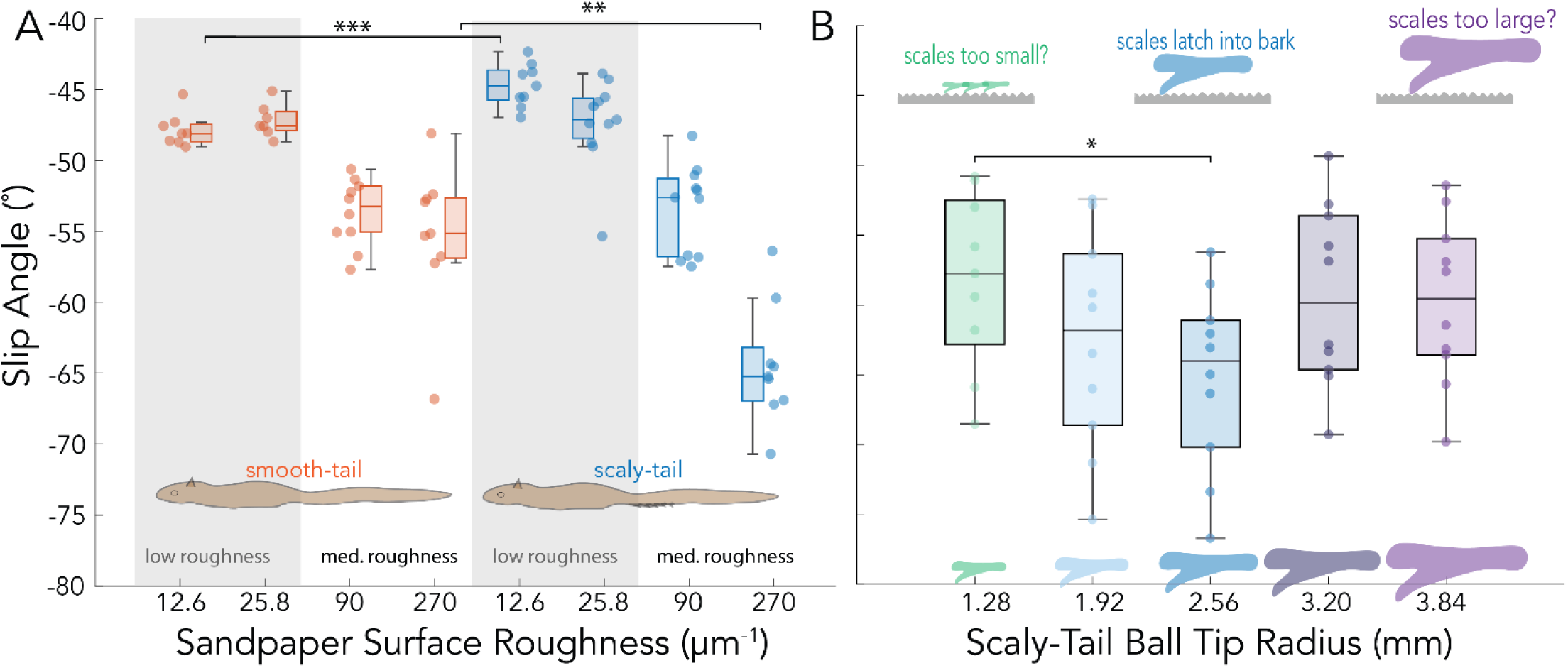
A) Display of the slip angle β of the smooth tail (orange) and scaly tail (blue) for four different surface roughness profiles of sandpaper ranging from 12.6 to 270.1. B) Slip angle (β) for five different scaly-tailed organ ball tip radii of curvature with insets that display the spines’ intermediate, large, and small values. Statistical significance shown with starts (^∗^ *p <* 0.05, ^∗∗^ *p <* 0.01, ^∗∗∗^ *p <* 0.001).

## 3. Results

### 3.1 Morphological Data

Morphological data of the Pel’s scaly-tailed squirrel specimen, shown in **Figure 3**A, included gross morphological measurements of the body, claw geometry (**Figure 3**B-C), and spiny tail (**Figure 3**D). The squirrel’s body length is 37.2 cm (**Figure 2**B). The tail length is 25.8 cm. The tail begins with a short hairy portion, 1.7 cm long, which is the vertical distance between the hind limb and the start of the tail’s scaly organ. The scaly organ is 6.9 cm long, approximately one-quarter of the tail length, similar to that of *A. derbianus* [22]. There are two rows of seven spines along the left and right sides of the scaly-tail organ, with bare skin present between each duo of scales (**Figure S2**). The scales’ size, area, and height gradually decrease distally towards the tip of the tail (**Figure 3**A,D). The length and width of the scales decrease distally from 7.5 x 8.5 mm to 3.2 x 3.3 mm; the length and width remain similar for all the scales. The area of the scales varies from 47.3 mm^2^ at the proximal end to 5.6 mm^2^ at the distal end. The scale height decreases distally from 6.4 mm at the proximal end to 2.6 mm at the distal end. For their limbs, the Pel’s scaly-tailed squirrel has four digits on its forelimbs and five digits on its hind limbs.

In comparing the three friction-enhancing structures, we see significant differences in the curvature of the point of the foreclaws, hindclaws, and caudal organs. Using Equation (1), we find the curvature in *mm*^−1^ to be 0.71, 0.54, and 0.39 for the foreclaw tip, hindclaw tip, and scaly spine tip, respectively (**Figure 3**B-D). The exact 3D structures of these claws and the scaly-tail organ were scanned and printed (**Figure 3**E), and all additional morphological and 3D scanned files can be found in the supplementary material. Using the squirrel’s scanned morphology, we mimicked each friction-enhancing structure and tested their specific contributions to friction enhancement on diverse substrates and their contributions toward sliding stability.

### 3.2 Sliding stability

#### 3.2.1 Substrate engagement and coefficient of static friction

The slip angle (β) for the squirrel model (with and without the scaly-tail organ) on each substrate is shown in **Figure 5**A along with the corresponding coefficient of static friction (*µ_s_*) in **Table 1**. The smooth tail model sustained ∼14% steeper perch angles (β = -54.4 ± 3.9^◦^, n = 19) on the medium roughness substrate than on the low roughness substrate (β = -47.5 ± 1.8^◦^, n = 15).The model with the scaly-tail organ performed poorly on the low roughness substrate with β even shallower than the smooth tail.

**Table 1:**
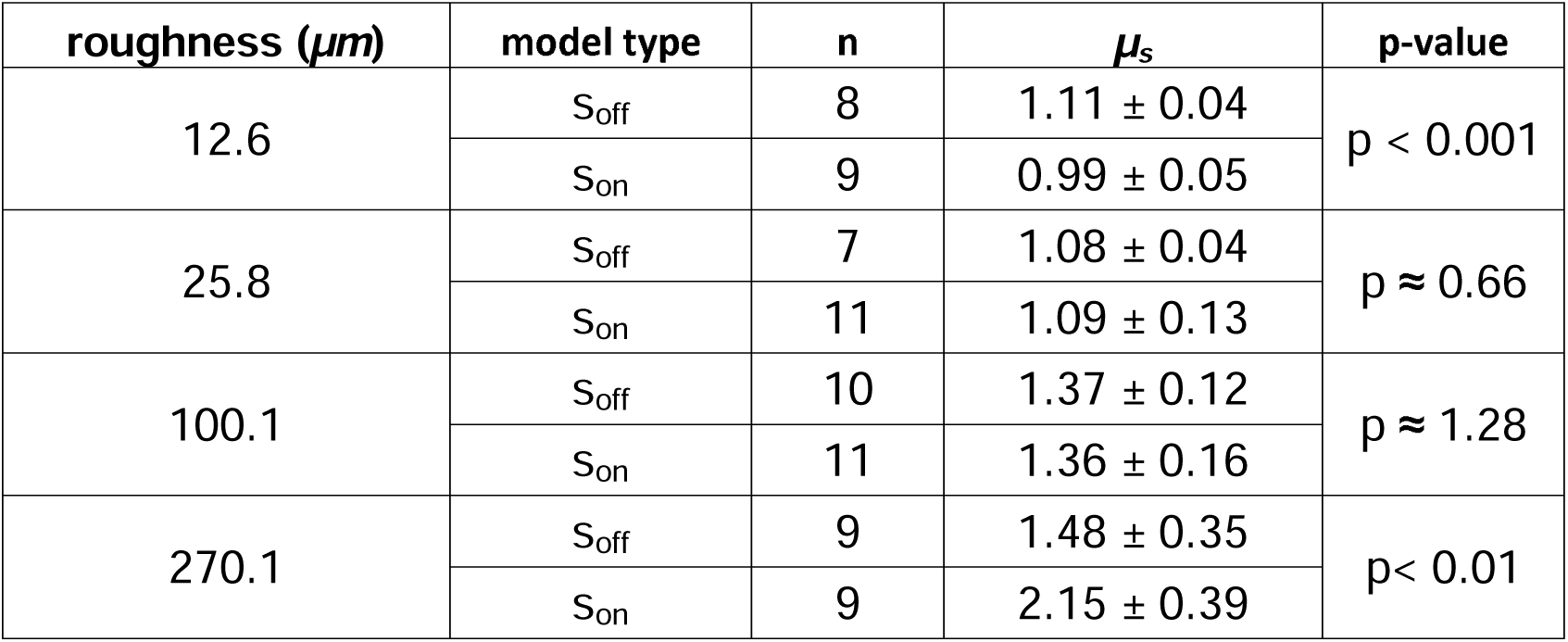
The coefficient of static friction *µ_s_* for each substrate with (*s_on_*) and without the scaly-tail organ (*s_off_*). The substrate with a surface roughness of 270.1 *µ*m showed a significant increase in substrate engagement with the scaly-tail organ.

The smooth and scaly-tail model had significantly higher substrate engagement on the medium roughness substrate compared to the low roughness substrate, as shown by the increase in the observed β and *µ_s_*. Furthermore, the substrate engagement of the scaly-tail model drastically increased on the 270.1 *µ*m roughness, resulting in a β of -64.5 ± 4.2^◦^ (n = 9) – a ∼21% jump in the perch steepness that could be sustained passively relative to the ∼90 *µ*m roughness substrate. Altogether, the maximum slip angle of the scaly-tail organ was seen on the 270.1 *µ*m substrate roughness, resulting in an increase of ∼17% in β along with a ∼58% increase in *µ_s_* of the squirrel model as a whole.

#### 3.2.2 Scaly-tail ball tip curvature

With a significant friction enhancement demonstrated with the scaly-tail organ on the 270.1 *µm*^−1^ substrate, we used the same substrate to test the influence of different scaly-tail organ sizes on substrate engagement by quantifying β and *µ_s_* (**Figure 5**B and **Table 2**). Five sizes of the scaly-tail organ were tested which included 0.5x (1.28 mm), 0.75x (1.92 mm), 1x (2.56 mm), 1.25x (3.20 mm), and 1.5x (3.84 mm). The mean slip angle varied between -58^◦^ and -65^◦^, with the steepest angle corresponding to the 1x scaled version of the organ (**Figure 5**B). The 1x version was significantly steeper than the .5x version, resulting in a ∼40% increase in mean *µ_s_* from 1.69 to 2.36 (**Table 2**).

**Table 2:** The coefficient of static friction *µ_s_* for different scaled versions of the organ. The 1x version showed the highest substrate engagement compared to the 0.5x, 0.75x, 1.25x, and 1.5x version.

| scale | n | $\mu_s$ | p-value |
| --- | --- | --- | --- |
| 0.5x | 9 | $1.69 \pm 0.46$ | **0.035** |
| 0.75x | 10 | $2.11 \pm 0.81$ | 0.473 |
| 1x | 10 | $2.36 \pm 1.28$ | – |
| 1.25x | 10 | $1.77 \pm 0.47$ | 0.104 |
| 1.5x | 10 | $1.79 \pm 0.46$ | 0.104 |

Though a U-shaped trend is indicated in **Figure 5**, the β and *µ_s_* values for the 1x version were not significantly different than the 0.75x, 1.25x, and 1.5x versions. The maximum slip angle of the scaly-tail organ was seen in the 1x version, resulting in an increase of 10% in β compared to the 1.25x and an increase of 3.5% compared to 0.75x.

### 3.3 Overturning stability

The presence of spiny-tail changes the shape and area of the support polygon. Moreover, when the scaly-tail organ’s length increases, the support polygon’s area further increases. When the scaly-tail organ’s length was increased by 100% from the original size (8.6 cm), the maximum inclination of the substrate the squirrel can perch without applying any force by the claws is increased by 2%. However, when the tail length was decreased by 50%, the maximum inclination angle was reduced by 1.5% (**Figure 6**A).

**Figure 6:**
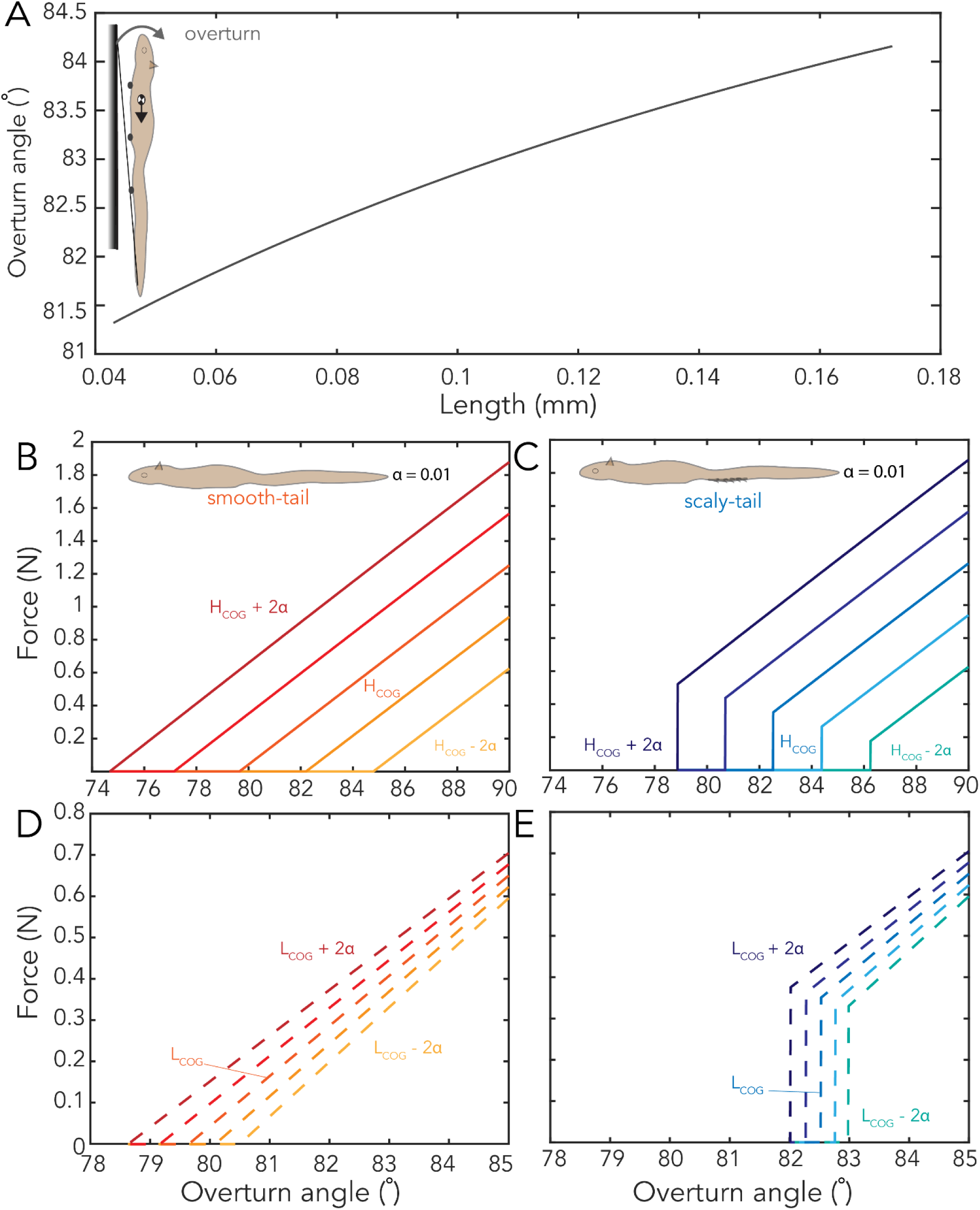
Overturning Stability. A) Variation of overturning angle for different lengths of spines and B-D) the force required by claws to counter the overturning for different locations of COG varying the height (B-C) and varying the length (D-E).

When the inclination increases beyond maximum inclination angle, the claws will have to actively apply additional force to prevent the overturn. From our model, a scaly-tailed squirrel with COG position at *H_COG_* = 4 cm and *L_COG_* = 17 cm can reach up to an inclination of 82.5^◦^ without applying any force; however, without the scales, the maximum inclination angle is reduced to 79.6^◦^ (**Figure 6**B-C).

We also analyzed the influence of the position of COG on the additional force required by the claws. **Figure 6**B-E shows the force needed for the claws to grab the substrate for different locations of COG. Our model also suggests that an increase in the height of COG results in an increase in the force required by the claws. For example, at a very steep inclination of 88^◦^, the force required by a hind claw increased from 0.39 N to 1.64 N when the height of the COG increased from 2 cm to 6 cm (**Figure 6**B-C). Similarly, at 84^◦^ of inclination, when the distance of COG from the snout increased from 15 cm to 19 cm, the force required by the claws increased from 0.46 N to 0.59 N (**Figure 6**D-E). The results show that the variation in the height of COG is more sensitive to the force required by the claws.

## 4. Discussion

The scaly-tail organ is a unique structure found at the tail base in the family *Anomaluridae*. The organ consists of multiple scales extruding from the tail bone, which can provide an additional mechanism to physically engage with the substrate along with the fore- and hindclaws. Based on this unique tail structure and the presence of ‘smoother’ bark trees in the scaly-tail squirrel’s habitat, we tested the hypothesis of the scaly-tail organ providing additional frictional enhancement (sliding stability) on substrates smoother than rough tree barks, and allowing the squirrel to sustain steeper perch angles without overturning (overturning stability).

Our results supported the hypothesis that the scaly-tail organ provided additional frictional benefits on intermediate rough tree barks but performed poorly on extremely smooth surfaces. Additionally, we found that the scales may be adapted to potentially maximize engagement on intermediate roughness surfaces that resemble the tree barks found in their natural habitat. Finally, we made a simple 2D squirrel model with the claws and the scaly-tail organ as contact points forming a support polygon. The 2D squirrel model showed that the scaly-tail organ can allow the squirrel to sustain steeper perch angles without actively engaging the claws.

### 4.1 Scales could be adapted for ‘smoother’ barked trees

The species examined in this study, Pel’s scaly-tailed flying squirrel, is geographically located in the southwest of Ghana and the southeast of Liberia. This is primarily located in the upper Guinea rainforest zone [41], which is a unique tropical zone with a large population of drought-tolerant tree species [42], [43], [44]. The drought-tolerant species such as *Milicia excelsa* have a smoother bark and are inhabited by the Pel’s scaly-tailed squirrel (**Figure 1**D). Previously, it has been hypothesized that the scaly-tail organ could be an adaptation of the squirrel’s arboreal lifestyle on trees, but no previous papers have connected the scales to smoother bark trees found in their drought-adapted rainforest habitat. Our results show that the scaly-tail organ can enhance the static frictional force on an intermediate roughness substrate that falls between a bamboo-like extremely smooth substrate and high roughness tree barks (**Figure 5**A). For the squirrel, this frictional enhancement could lead to less slipping and reduce the energetic cost of actively engaging with the substrate for elongated periods of perching. However, no frictional enhancement was observed on smooth substrates where the squirrel physical model with and without the scaly-tail organ resulted in similar slip angles, suggesting that the scale size was likely too large to engage with the smooth substrate.

Varying the scaly-tail organ size showed that the actual scale size (1x version) had the maximum engagement, resulting in a ∼40% increase in mean *µ_s_* from 1.69 to 2.36 compared to the scaled-down 0.5x version. It should be noted that the 1x version did not show a significant difference compared to the 0.75x, 1.25x, and 1.5x version. Nonetheless, the significant enhancement in static friction on the intermediate roughness substrate combined with the U-shaped slip angle trend from the 0.5x to the 1.5x version (**Figure 5**B), suggest that the scaly-tail organ could be an adaptation to enhance the perching capabilities of the scaly-tailed squirrel on the ‘smooth’ barks that are found in their natural habitat. By enhancing the interaction with the arboreal substrate, adaptations such as the scaly-tail organ can provide instantaneous mechanical feedback and compensate for the loss in frictional forces from unforeseen perturbations experienced by the animal.

Comparing the limited morphology data available on these elusive scaly-tailed squirrels, we see that the Pel’s scaly-tailed squirrel had the longest scaly tail organ of 69 mm compared to *A. beecrofti*, *A. derbianus*, and *A. pusillus* that had lengths of 40, 64, and 34 mm, respectively [16]. It is possible that as the Pel’s scaly-tailed squirrel has the longest scaly tail organ, this allows the most robust attachment as in other species like geckos, larger size and area allows for higher maximum force [45]. Future studies should compare the scaly-tail organ engagement across species to identify if the organ size and shape are adapted to the tree type or vary based on the species body size.

### 4.2 Contribution of scaly-tail organ towards overturning stability

The presence of friction enhancement features to scale arboreal substrates are common in the animal kingdom [6]. Numerous studies on lizards have shown the advantages of claws for clinging and interacting with various substrates [8], [46], [47]. Interestingly, the scaly-tail organ not only provides frictional enhancements similar to the fore- and hindclaw, but its anatomical location can act as a fifth point of contact, enlarging the area within which the COG lies during perching. This fifth point of contact could help offset the fact that larger animals often attach less well to surfaces [6]. The area formed, referred to as a support polygon, can allow the squirrel to maintain passive stability while perching without risking overturning and falling to the ground. Our 2D mathematical model for overturning stability showed that the scaly-tail organ can enable the squirrel to sustain ∼a 4% steeper perch angle passively compared to a four-point support polygon formed by the fore- and hindclaws only. Since the steeper perch angle is a consequence of passive perching, this can significantly reduce the energetic costs of perching for the scaly-tailed squirrel on ‘smooth’ tree barks without applying additional force through their claws to prevent overturning.

### 4.3 Bio-inspired Friction Enhancement for Perching

Arboreal environments are challenging to explore and collect data [48], [49]. Robotics engineers have focused on two strategies to explore arboreal environments: climbing robots and aerial robots [50]. In both these approaches, the robots often must perch on the tree trunks and branches to collect the environmental data. Perching allows the robots to depend less on their locomotion modes to stay stationary, resulting in lesser control effort and more battery life. Therefore, perching extends the total mission time of the robot during their operation in challenging environments like the thick canopies of arboreal environments. Various methods have been introduced so far to address the perching challenges, such as gecko-inspired adhesives, mechanical spines, and bird-inspired claws [51], [52], [53], [54].

Our study on the scaly-tailed squirrel’s caudal scaly-tail organ provides valuable insights for innovative robot perching mechanisms. By incorporating these insights, we can introduce a novel design that enhances the sliding and overturning stability of the robot during perching. One approach involves integrating additional support points through passive spines on the chassis of the robot. These spines mimic the functionality of the squirrel’s caudal scaly-tail organ, enabling the robot to securely grip onto branches and tree trunks. Spines enhance the sliding stability by increasing the frictional resistance and improve overturning stability by increasing the area of the support polygon, as the spines act as an additional support point.

Additionally, the implementation of spines on additional appendages with morphing capabilities [55], [56] could offer further versatility in perching. These appendages can adapt to varying surfaces and angles, providing the robot with enhanced stability on various surfaces. Furthermore, customizing the morphology of the scales to match the nature of the trees in the arboreal environment where the robot operates is also crucial. Potentially optimizing the angle of the claws could have impacts assisting with friction enhancement as it has been modeled in small vertebrates [57]. By selecting scale structures tailored to the roughness of the bark, the robot can optimize its stability, thereby improving overall performance and mission success in challenging arboreal environments.

### 4.4 Study Limitations

Scaly-tailed squirrels are one of the least studied mammalian clades [22], which could be because of their arboreal and nocturnal lifestyle and their presence in a small geographical region. With how understudied these species are, we had limited access to museum specimens, limiting our morphological data to a single mounted skin. This limitation of single-mounted skin also did not allow us to compare the hindlimb to the forelimb for muscle density. Therefore, we used the comparison of limb length to estimate force differences between the fore and hind limbs.

Moreover, we could not observe live animals in the wild and do not know to what degree scaly-tailed squirrels actively control their compliant tail. Therefore, we assumed the tail and body are rigidly connected. This assumption is justified as our study focuses only on the role of spiny-tailed organs on the perching stability of the squirrel. The role of compliance of the body and the tail in the perching stability is an important area that can be explored in future studies.

In our experiments, we could not do mechanical frictional testing on any native trees for Pel’s scaly-tailed squirrel. In future studies of the scaly-tailed squirrel, it would be worthwhile to benchmark the frictional surfaces they often interact with, which has been done in many North American gliding squirrel species[36]. Additionally, the frictional comparison of bark to sandpaper was not within the scope of this study and was based on inferences drawn from a previous study [35]. In our trials, our scaly-tail model trials followed the smooth-tail trials, potentially meaning the poor performance of the scaly-tail model could be an artifact of the claws degrading, leading to lower substrate engagement. Future studies should examine and incorporate the frictional properties of different rough to smooth-like barks to better understand the interaction between the scaly-tail organ and the substrate.

## 5. Conclusion

Our study provides novel insights into the scaly-tail organ’s contribution towards enhancing the stability of the scaly-tailed flying squirrel in its natural habitat. Our results suggest that the scaly-tail organ could be an adaptation for the squirrel to increase friction and reduce energetic costs while perching in their ‘smooth’ bark tree habitat. Using printed mimics from the 3D scans, we demonstrated that the scaly-tailed organ could help reduce skid and provide significant frictional benefits on surfaces of similar roughness to trees in its natural habitat. Additionally, our findings show that the scale size is likely adapted to maximize engagement with intermediate roughness substrates. Finally, through simulation, we showed that the scaly-tail organ enhances the squirrel’s overturning stability by acting as an additional contact point.

Our findings highlight the importance of using bioinspired models to understand the complex mechanics of climbing adaptations in species that are challenging to study. To the best of our knowledge, this is the first study on these unique arboreal rodents that provides support for the role of the scaly-tail organ to enhance perching stability, especially in the context of their natural arboreal environment. Altogether, our study provides a deeper understanding of the environmental stresses that might have driven the adaptation of the scaly-tail organ and identifies aspects of the tail morphology that could be translated to robot design to improve their locomotion capability on incline and vertical substrates.

## Data Access Statement

All data will be accessible in an Edmond repository of MPG [58]. The museum specimen’s body measurements and scaly-tail organ measurements are included in the supplementary file.

## Ethics Statement

Measurements and scans were taken from the mounted skin SMNS-Z-MAM-001377 preserved at the Natural History Museum Stuttgart.

## Author Contributions

Conceptualization: A.S. and A.J., Data curation: A.S., S.M., P.K., M.C., A.J., Formal Analysis: A.S., S.M., P.K., M.C., Funding Acquisition: A.J. Investigation: A.S., P.K., M.C., R.R., A.J. Methodology: A.S., P.K., M.C., R.R., A.J., Project administration: A.S., P.K., M.C., Resources: A.J. Software: P.K., M.C., R.R., Supervision: A.S., Validation: A.S., P.K., M.C., R.R., Visualization: A.S., P.K., M.C., Writing – Original Draft: A.S., P.K., M.C., R.R., Writing - review & editing: A.S., P.K., M.C., R.R., S.M., A.J. Finally, A.S., M.C., P.K. contributed equally and have the right to list their first name first in their CV.

## Acknowledgements

A.S. thanks the support of the International Max Planck Research School for Intelligent Systems (IMPRS-IS) in Stuttgart, Germany. A.J. thanks the Swiss National Science Foundation for grants in support of research. The authors thank Marc and Peggy Faucher for their HD photo of Pel’s scaly-tailed squirrel for this manuscript and other images that can be found at (https://www.mfaucher.com/). We thank the Central Service Station for Robotics at Max Planck Institute for Intelligent Systems for a loan of a handheld 3d scanner and for assisting with the 3d printing of scaly-tail squirrel’s claws and tail spines. We thank the State Museum of Natural History Stuttgart for specimen access.

## Bibliography

[1] H. Preuschoft, “What does ‘arboreal locomotion’ mean exactly and what are the relationships between ‘climbing’, environment and morphology?,” Zeitschrift für Morphologie und Anthropologie, vol. 83, no. 2/3, pp. 171–188, 2002.

[2] J. W. Young, “Convergence of Arboreal Locomotor Specialization: Morphological and Behavioral Solutions for Movement on Narrow and Compliant Supports,” in *Convergent Evolution: Animal Form and Function*, V. L. Bels and A. P. Russell, Eds., Cham: Springer International Publishing, 2023, pp. 289–322. doi: 10.1007/978-3-031-11441-0_11.

[3] M. Hildebrand, D. M. Bramble, K. F. Liem, and D. B. Wake, Functional Vertebrate Morphology. Harvard University Press, 2013.

[4] G. Byrnes and B. C. Jayne, “Gripping during climbing of arboreal snakes may be safe but not economical,” Biol Lett, vol. 10, no. 8, p. 20140434, Aug. 2014, doi: 10.1098/rsbl.2014.0434.

[5] A. Biewener and S. Patek, Animal Locomotion, Second Edition. Oxford, New York: Oxford University Press, 2018.

[6] D. Labonte and W. Federle, “Scaling and biomechanics of surface attachment in climbing animals,” Philosophical Transactions of the Royal Society B: Biological Sciences, vol. 370, no. 1661, p. 20140027, Feb. 2015, doi: 10.1098/rstb.2014.0027.

[7] A. P. Russell, “A contribution to the functional analysis of the foot of the Tokay, Gekko gecko (Reptilia: Gekkonidae),” Journal of Zoology, vol. 176, no. 4, pp. 437–476, 1975, doi: 10.1111/j.1469-7998.1975.tb03215.x.

[8] N. Bloch and D. J. Irschick, “Toe-Clipping Dramatically Reduces Clinging Performance in a Pad-Bearing Lizard (Anolis carolinensis),” hpet, vol. 39, no. 2, pp. 288–293, Jun. 2005, doi: 10.1670/97-04N.

[9] T. E. Higham, “Lizard Locomotion: Relationships between Behavior, Performance, and Function,” in Behavior of Lizards, CRC Press, 2019.

[10] D. Santos, B. Heyneman, S. Kim, N. Esparza, and M. R. Cutkosky, “Gecko-inspired climbing behaviors on vertical and overhanging surfaces,” in 2008 IEEE International Conference on Robotics and Automation, May 2008, pp. 1125–1131. doi: 10.1109/ROBOT.2008.4543355.

[11] N. Bloch and D. J. Irschick, “Toe-Clipping Dramatically Reduces Clinging Performance in a Pad-Bearing Lizard (Anolis carolinensis),” Journal of Herpetology, vol. 39, no. 2, pp. 288–293, 2005.

[12] C. Krause and M. S. Fischer, “Biodynamics of climbing: effects of substrate orientation on the locomotion of a highly arboreal lizard (Chamaeleo calyptratus),” Journal of Experimental Biology, vol. 216, no. 8, pp. 1448–1457, Apr. 2013, doi: 10.1242/jeb.082586.

[13] E. Dickinson et al., “Tail feather strength in tail-assisted climbing birds is achieved through geometric, not material change,” Proceedings of the Royal Society B: Biological Sciences, vol. 290, no. 1998, p. 20222325, May 2023, doi: 10.1098/rspb.2022.2325.

[14] R. Å. Norberg, “Treecreeper climbing; mechanics, energetics, and structural adaptations,” Ornis scandinavica, vol. 17, no. 3, pp. 191–209, Jul. 1986, doi: 10.2307/3676828.

[15] A. Jusufi, D. I. Goldman, S. Revzen, and R. J. Full, “Active tails enhance arboreal acrobatics in geckos,” Proceedings of the National Academy of Sciences, vol. 105, no. 11, pp. 4215–4219, Mar. 2008, doi: 10.1073/pnas.0711944105.

[16] A. Jusufi, “The role of the tail in stability and maneuverability during running, climbing, mid-air orientation and gliding in both animals and robots.,” UC Berkeley, 2013. Accessed: Sep. 26, 2024. [Online]. Available: https://escholarship.org/uc/item/2hn391sc

[17] R. Siddall, G. Byrnes, R. J. Full, and A. Jusufi, “Tails stabilize landing of gliding geckos crashing head-first into tree trunks,” Commun Biol, vol. 4, no. 1, p. 1020, Sep. 2021, doi: 10.1038/s42003-021-02378-6.

[18] A. Jusufi, D. T. Kawano, T. Libby, and R. J. Full, “Righting and turning in mid-air using appendage inertia: reptile tails, analytical models and bio-inspired robots,” Bioinspir. Biomim., vol. 5, no. 4, p. 045001, Nov. 2010, doi: 10.1088/1748-3182/5/4/045001.

[19] T. Libby et al., “Tail-assisted pitch control in lizards, robots and dinosaurs,” Nature, vol. 481, no. 7380, pp. 181–184, Jan. 2012, doi: 10.1038/nature10710.

[20] T. Fukushima et al., “Inertial Tail Effects during Righting of Squirrels in Unexpected Falls: From Behavior to Robotics,” Integrative and Comparative Biology, vol. 61, no. 2, pp. 589–602, Aug. 2021, doi: 10.1093/icb/icab023.

[21] M. Chellapurath, P. Khandelwal, T. Rottier, F. Schwab, and A. Jusufi, “Morphologically Adaptive Crash Landing on a Wall: Soft-Bodied Models of Gliding Geckos with Varying Material Stiffnesses,” Advanced Intelligent Systems, vol. n/a, no. n/a, p. 2200120, doi: 10.1002/aisy.202200120.

[22] A. A. Panyutina, O. F. Chernova, Irina Soldatova, and I. Soldatova, “Morphological peculiarities in the integument of enigmatic anomalurid gliders (Anomaluridae, Rodentia).,” Journal of Anatomy, vol. 237, no. 3, pp. 404–426, May 2020, doi: 10.1111/joa.13211.

[23] V. Dinets, “First observations on the behavior of the flightless anomalure (*Zenkerella insignis*),” Zoology, vol. 123, pp. 121–123, Aug. 2017, doi: 10.1016/j.zool.2017.06.003.

[24] V. Nicolai, “Thermal properties and fauna on the bark of trees in two different African ecosystems,” Oecologia, vol. 80, no. 3, pp. 421–430, Aug. 1989, doi: 10.1007/BF00379046.

[25] IUCN/SSC, Guidelines for reintroductions and other conservation translocations. Version 1.0. IUCN Species Survival Commission Gland, Switzerland, 2013.

[26] S. Heritage, D. Fernández, H. M. Sallam, D. T. Cronin, J. M. Esara Echube, and E. R. Seiffert, “Ancient phylogenetic divergence of the enigmatic African rodent Zenkerella and the origin of anomalurid gliding,” PeerJ, vol. 4, p. e2320, 2016, doi: 10.7717/peerj.2320.

[27] D. R. Rosevear, The rodents of west Africa. Trustees of the British Museum (Natural History), 1969.

[28] R. M. Nowak, Walker’s mammals of the world. Baltimore: Johns Hopkins University Press, 1999.

[29] J. Kingdon, Mammals of Africa: Volume III: Rodents, Hares and Rabbits. A&C Black, 2014.

[30] S. M. Jackson and R. W. Thorington, “Gliding Mammals: Taxonomy of Living and Extinct Species,” no. 638, pp. 1–117, Jan. 2012, doi: 10.5479/si.00810282.638.1.

[31] C. Jones, “NOTES ON THE ANOMALURIDS OF RIO MUNI AND ADJACENT AREAS,” Journal of Mammalogy, vol. 52, no. 3, pp. 568–572, Aug. 1971, doi: 10.2307/1378591.

[32] P.-B. Wieber, R. Tedrake, and S. Kuindersma, “Modeling and Control of Legged Robots,” in *Springer Handbook of Robotics*, B. Siciliano and O. Khatib, Eds., Cham: Springer International Publishing, 2016, pp. 1203–1234. doi: 10.1007/978-3-319-32552-1_48.

[33] stratasys Ltd., “VeroClear,” Datasheet MDS_PJ_Vero_1020a, 2020.

[34] K. H. Orvis and H. Grissino-Mayer, “Standardizing the Reporting of Abrasive Papers Used to Surface Tree-Ring Samples,” Tree-Ring Society, vol. 58, no. 47–50, 2002.

[35] R. Pillai, E. Nordberg, J. Riedel, and L. Schwarzkopf, “Nonlinear variation in clinging performance with surface roughness in geckos,” Ecology and Evolution, vol. 10, no. 5, pp. 2597–2607, 2020, doi: 10.1002/ece3.6090.

[36] C. A. Diggins, J. L. D. L. Cruz, and A. Silvis, “Distribution Probability of the Virginia Northern Flying Squirrel in the High Allegheny Mountains,” Southeastern Association of Fish and Wildlife Agencies, Mar. 2022. Accessed: Apr. 27, 2024. [Online]. Available: https://seafwa.org/journal/2022/distribution-probability-virginia-northern-flying-squirrel-high-allegheny-mountains

[37] A. Mathis et al., “DeepLabCut: markerless pose estimation of user-defined body parts with deep learning,” Nat Neurosci, vol. 21, no. 9, Art. no. 9, Sep. 2018, doi: 10.1038/s41593-018-0209-y.

[38] M. H. Raibert, Legged Robots that Balance. MIT Press, 1986.

[39] R. W. Thorington Jr. and L. R. Heaney, “Body Proportions and Gliding Adaptations of Flying Squirrels (Petauristinae),” Journal of Mammalogy, vol. 62, no. 1, pp. 101–114, Mar. 1981, doi: 10.2307/1380481.

[40] B. M. Kilbourne and L. C. Hoffman, “Scale Effects between Body Size and Limb Design in Quadrupedal Mammals,” PLoS ONE, vol. 8, no. 11, p. e78392, Nov. 2013, doi: 10.1371/journal.pone.0078392.

[41] B. Y. Ofori and D. K. Attuquayefio, “Hunting Intensity in the Suhuma Forest Reserve in the Sefwi Wiawso District of the Western Region of Ghana: A Threat to Biodiversity Conservation,” West African Journal of Applied Ecology, vol. 17, no. 1, Art. no. 1, 2010, doi: 10.4314/wajae.v17i1.65142.

[42] S. Asefi-Najafabady and S. Saatchi, “Response of African humid tropical forests to recent rainfall anomalies,” Philos Trans R Soc Lond B Biol Sci, vol. 368, no. 1625, p. 20120306, 2013, doi: 10.1098/rstb.2012.0306.

[43] Y. Malhi, S. Adu-Bredu, R. A. Asare, S. L. Lewis, and P. Mayaux, “African rainforests: past, present and future,” Philos Trans R Soc Lond B Biol Sci, vol. 368, no. 1625, p. 20120312, 2013, doi: 10.1098/rstb.2012.0312.

[44] S. Fauset et al., “Drought-induced shifts in the floristic and functional composition of tropical forests in Ghana,” Ecol Lett, vol. 15, no. 10, pp. 1120–1129, Oct. 2012, doi: 10.1111/j.1461-0248.2012.01834.x.

[45] C. A. Gilman, M. J. Imburgia, M. D. Bartlett, D. R. King, A. J. Crosby, and D. J. Irschick, “Geckos as Springs: Mechanics Explain Across-Species Scaling of Adhesion,” PLoS One, vol. 10, no. 9, p. e0134604, Sep. 2015, doi: 10.1371/journal.pone.0134604.

[46] D. C. D’Amore, S. Clulow, J. S. Doody, D. Rhind, and C. R. McHenry, “Claw morphometrics in monitor lizards: Variable substrate and habitat use correlate to shape diversity within a predator guild,” Ecology and Evolution, vol. 8, no. 13, pp. 6766–6778, 2018, doi: 10.1002/ece3.4185.

[47] J. C. Spagna, D. I. Goldman, P.-C. Lin, D. E. Koditschek, and R. J. Full, “Distributed mechanical feedback in arthropods and robots simplifies control of rapid running on challenging terrain,” Bioinspir. Biomim., vol. 2, no. 1, p. 9, Jan. 2007, doi: 10.1088/1748-3182/2/1/002.

[48] M. Chellapurath, P. C. Khandelwal, and A. K. Schulz, “Bioinspired robots can foster nature conservation,” Front. Robot. AI, vol. 10, Oct. 2023, doi: 10.3389/frobt.2023.1145798.

[49] S. Kirchgeorg and S. Mintchev, “HEDGEHOG: Drone Perching on Tree Branches With High-Friction Origami Spines,” IEEE Robotics and Automation Letters, vol. 7, no. 1, pp. 602–609, Jan. 2022, doi: 10.1109/LRA.2021.3130378.

[50] S. Kirchgeorg, E. Aucone, F. Wenk, and S. Mintchev, “Design, Modeling, and Control of AVOCADO: A Multimodal Aerial-Tethered Robot for Tree Canopy Exploration,” IEEE Transactions on Robotics, vol. 40, pp. 592–605, 2024, doi: 10.1109/TRO.2023.3334630.

[51] K. Autumn et al., “Adhesive force of a single gecko foot-hair,” Nature, vol. 405, no. 6787, pp. 681–685, Jun. 2000, doi: 10.1038/35015073.

[52] G. Byrnes and A. J. Spence, “Ecological and biomechanical insights into the evolution of gliding in mammals,” Integr Comp Biol, vol. 51, no. 6, pp. 991–1001, Dec. 2011, doi: 10.1093/icb/icr069.

[53] E. W. Hawkes, E. V. Eason, A. T. Asbeck, and M. R. Cutkosky, “The Gecko’s Toe: Scaling Directional Adhesives for Climbing Applications,” IEEE/ASME Transactions on Mechatronics, vol. 18, no. 2, pp. 518– 526, Apr. 2013, doi: 10.1109/TMECH.2012.2209672.

[54] M. Kova, J. Germann, C. Hürzeler, R. Y. Siegwart, and D. Floreano, “A perching mechanism for micro aerial ve^č^hicles,” J. Micro-Nano Mech., vol. 5, no. 3, pp. 77–91, Dec. 2009, doi: 10.1007/s12213-010-0026-1.

[55] P. Zheng, F. Xiao, P. H. Nguyen, A. Farinha, and M. Kovac, “Metamorphic aerial robot capable of mid-air shape morphing for rapid perching,” Sci Rep, vol. 13, no. 1, p. 1297, Jan. 2023, doi: 10.1038/s41598-022-26066-5.

[56] J. Meng, J. Buzzatto, Y. Liu, and M. Liarokapis, “On Aerial Robots with Grasping and Perching Capabilities: A Comprehensive Review,” Front. Robot. AI, vol. 8, Mar. 2022, doi: 10.3389/frobt.2021.739173.

[57] D. Labonte and W. Federle, “Biomechanics of shear-sensitive adhesion in climbing animals: peeling, pre-tension and sliding-induced changes in interface strength,” Journal of The Royal Society Interface, vol. 13, no. 122, p. 20160373, Sep. 2016, doi: 10.1098/rsif.2016.0373.

[58] A. K. Schulz, M. Chellapurath, P. Khandelwal, S. Rezaei, S. Merker, and A. Jusufi, “Data for: Scaly-Tail Organ Enhances Static Stability During the Perching of Pel’s Scaly-tailed Flying Squirrel.” Edmond. doi: 10.17617/3.HAV0R2.

