## Supplemental Tables and Figures for "Scaly-Tail Organ Enhances Static Stability during Pel’s Scaly-tailed Flying Squirrels’ Arboreal Locomotion"

A. K. Schulz, M. Chellapurath, and P. Khandelwal et al.

Supplementary Tables:

**Table S1:** Table displaying the diversity of the phylogeny of the *Rodentia* order, including the Anomaluridae family, which we examined in this paper, as well as their distant relatives, including tree squirrels as well as flying and gliding squirrels in different families.

| Order | Family | Genus | Species Common Name |
| --- | --- | --- | --- |
| *R*odentia | Anomaluridae | *Anomalurus* | *pelii* Pel’s scaly-tailed squirrel |
| *R*odentia | Anomaluridae | *Anomalurus* | *beecrofti* Beecroft’s scaly-tailed squirrel |
| *R*odentia | Anomaluridae | *Anomalurus* | *derbianus* Lord Derby’s scaly-tailed squirrel |
| *R*odentia | Anomaluridae | *Anomalurus* | *pusillus* dwarf scaly-tailed squirrel |
| *R*odentia | Pedetidae | *Pedetes* | *capensis* South African springhare |
| *R*odentia | Sciuridae | *Glaucomys* | *sabrinus* Northern flying squirrel |
| *R*odentia | Sciuridae | *Sciurus* | *vulgaris* Red Squirrel |

**Table S2:** Table displaying the length measurements of the museum specimen.

| Length Of | Variable | Length (cm) |
| --- | --- | --- |
| Head | *Lhead* | 10.2 |
| Neck | *Lneck* | 1.7 |
| Forelimb | *Lforelimb* | 20.3 |
| Torso | *Ltorso* | 27.0 |
| Body (excluding tail) | *L_B_* | 37.2 |
| Hindlimb | *Lhindlimb* | 12.1 |
| Proximal tail without spines | *Ltail,pre* | 1.76 |
| Length of organ | *Ltail,organ* | 6.86 |
| Distal tail without spines | *Ltail,post* | 17.2 |
| Total Tail Length | *Ltail* | 25.82 |
| Width of head | *Dhead* | 4.3 |
| Forelimb width | *Dforelimb* | 3.2 |
| Center of torso width | *Dtorso* | 7.2 |
| Hindlimb width | *Dhindlimb* | 3.77 |
| Proximal tail width | *Dtail,*0 | 2.77 |
| distal tail tip width | *Dtail,f* | 2.25 |

**Table S3:** Displays the total experimental tests performed and what substrates were tested with what spiny and clawed appendage combinations.

| Foreclaws | Hindclaws | Tail Substrate | Surface Substate | Surface Roughness (*µ*m) | Trials |
| --- | --- | --- | --- | --- | --- |
| Yes | Yes | plexiglass | 1 | 0 | 10 |
| Yes | Yes | plexiglass | 2 | 12.6 | 10 |
| Yes | Yes | plexiglass | 3 | 25.8 | 10 |
| Yes | Yes | plexiglass | 4 | 90 | 10 |
| Yes | Yes | plexiglass | 5 | 270 | 10 |
| Yes | Yes | Spines | 1 | 0 | 10 |
| Yes | Yes | Spines | 2 | 12.6 | 10 |
| Yes | Yes | Spines | 3 | 25.8 | 10 |
| Yes | Yes | Spines | 4 | 90 | 10 |
| Yes | Yes | Spines | 5 | 270 | 10 |

Supplementary Figures:

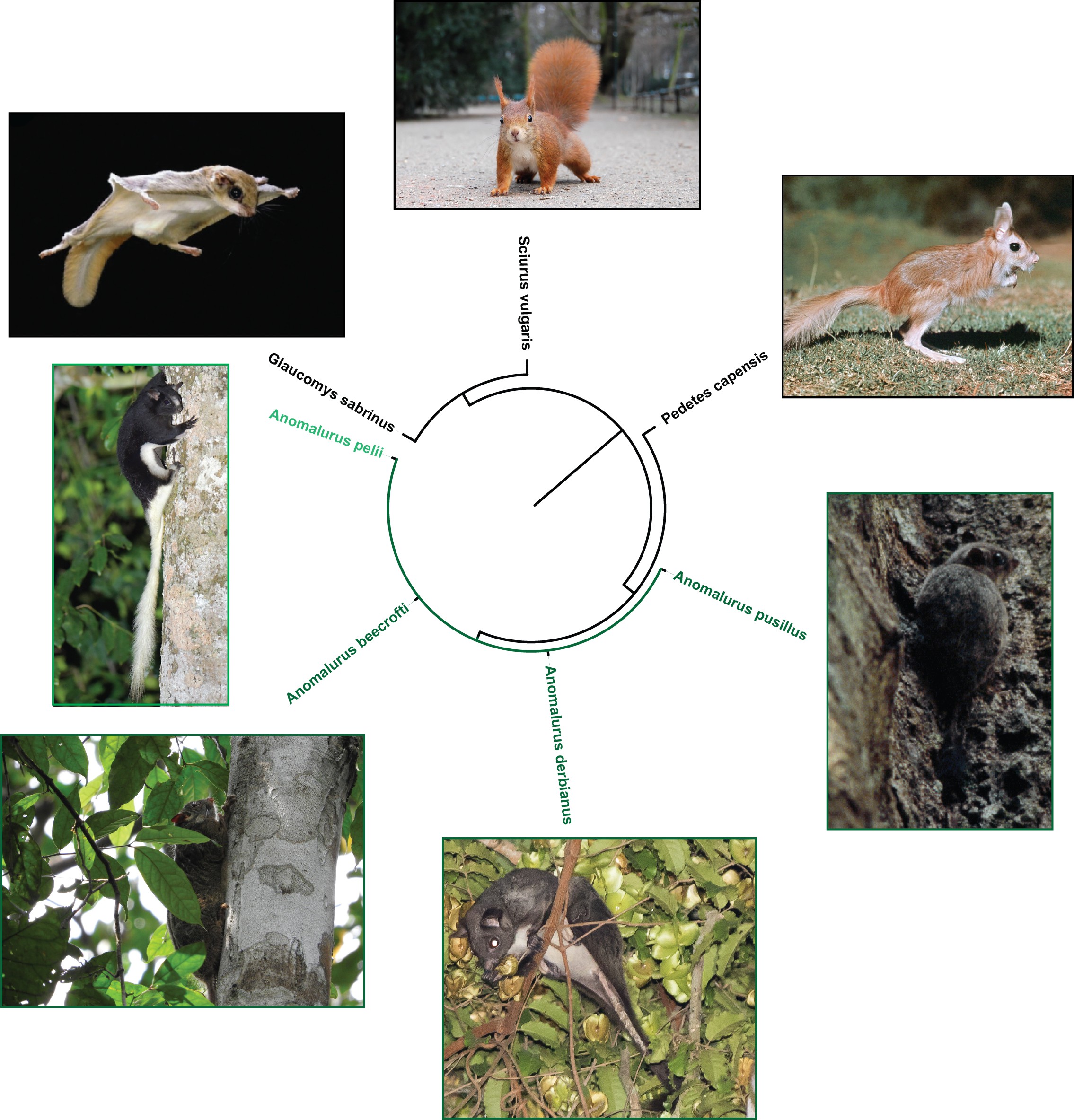

**Figure S1:** Phylogeny of rodents including the species of family Anomaluridae, or anomalures highlighted in green, including Dwarf scaly-tailed squirrel (*Anomalurus pusillus* taken by galat-luong-anh, Beecroft’s scaly-tailed squirrel (*Anomalurus beecrofti*) taken by pfaucher, Lord Derby’s scaly- tailed squirrel (*Anomalurus derbianus*) taken by jonhall100, and finally the species studied in this paper (in light green) Pel’s scaly-tailed squirrel (*Anomalurus pelii* ) taken by charleyhesse. Photos of springhare, northern flying squirrel, and red squirrel are also included. Details of each species and common names are included in Table S1. Photos are taken from iNaturalist, all under creative license CC BY-NC 4.0.

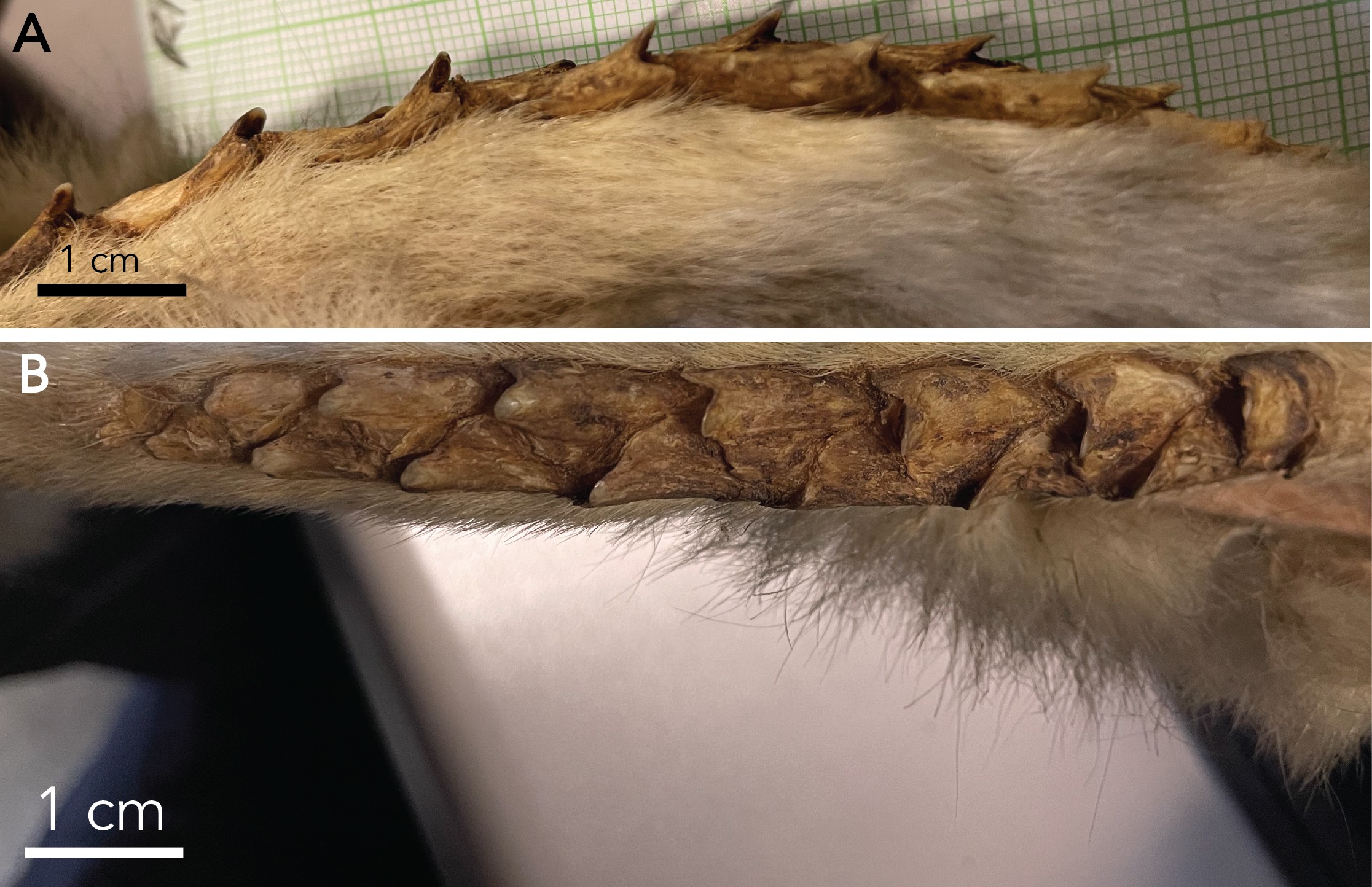

**Figure S2:** Images were taken of the specimen (SMNS-Z-MAM-001377), both A) side view and B) top- down view of the scaly organ of the squirrel.

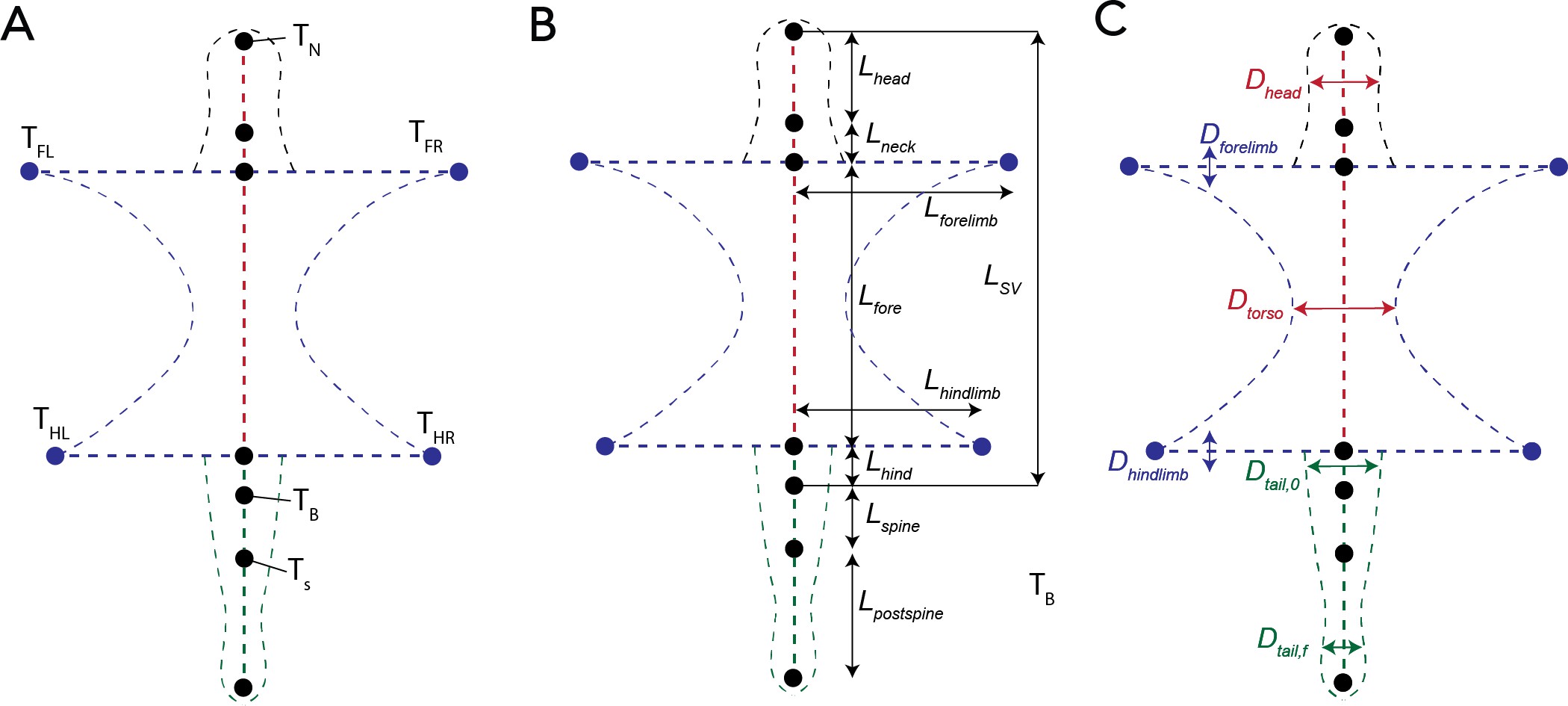

**Figure S3:** Schematic of different parameters used to describe the morphology of the squirrel museum specimen, including: A) Points along the squirrel’s body used for distance and force inputs. B) Length distances along the squirrel body between points of interest. C) Diameters of different appendages of the squirrel body. Exact measurements and descriptions of all labels are included in Table S2.
